# The lipid kinase PI3Kα is required for the cohesion and survival of cancer cells disseminated in serous cavities

**DOI:** 10.1101/777649

**Authors:** B Thibault, A Thole, C Basset, J Guillermet-Guibert

**Author notes:** Equal contribution (B Thibault, A Thole). **corresponding author:** Julie Guillermet-Guibert, CRCT UMR1037 INSERM-Université Toulouse 3 - ERL5294 CNRS; 2 avenue Hubert, Curien; Oncopole de Toulouse; CS 53717; 31037 TOULOUSE CEDEX 1 - FRANCE. http://www.crct-inserm.fr/17-j-guillermet-guibert-sigdyn-group-pi3k-isoforms-signalling-cancerogenesis-559.html, http://eupancreas.com/julie-guillermet-guibert, http://pi3k-phdproject.eu/partner/julie-guillermet-guibert/.

## Abstract

Breast, ovarian, digestive and lung adenocarcinomas are often associated with the accumulation of malignant cells in serous cavities. As PI3K is one of the most mutated pathways in cancer, we investigated the importance of oncogenic PI3Kα in this process. We analyzed tumor cell organization in ascites from carcinomas at diagnosis. In some malignant ascites, tumor cells grew as adhesive coherent masses. Ex-vivo patient-derived cell cultures with the addition of mesenchymal stem cells, as a model of tumoral stroma, favored the compaction of tumorospheres. Ascites-derived ovarian cancer cell lines frequently harbored *PIK3CA* mutations coexisting with other mutations. PI3Kα promoted the formation and maintenance of multicellular adhesive *PIK3CA*-mutant spheroids, promoting cell survival. Cultures in 3D conditions as opposed to cultures in cell monolayers increased chemotherapy resistance, which was overcome by PI3Kα inhibition. We identified a signaling pathway of interest for the treatment of cancer cells disseminated in serous cavities, limiting cancer progression.

**Graphical abstract:** Schematic representation of PI3Kα involvement in tumor cell aggregates from ascites. 1) Known involvement of PI3Kα in primary ovarian tumors. 2) PI3Kα participates in tumorosphere formation within the peritoneum (treatment with PI3Kα inhibitors causes a delay in the formation of clusters). 3) PI3Kα participates in the maintenance of tumorospheres and in resistance to conventional treatment for peritoneal carcinomatosis. PI3Kα is a target to prevent transcoelomic dissemination and maintenance of tumorospheres in patients with ovarian cancer.

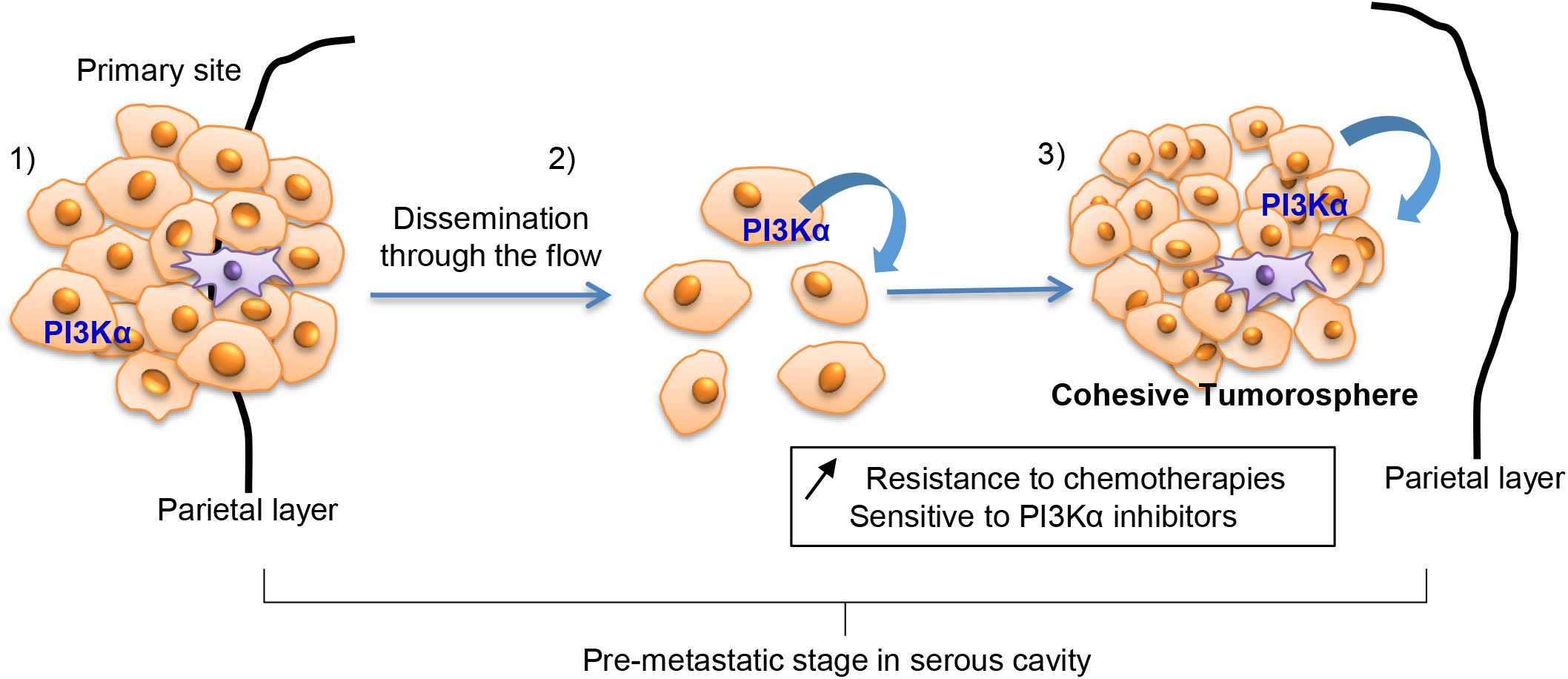

## Introduction

Tumor cell resistance to chemotherapy is now largely considered to be enhanced in multicellular 3D structures. For example, EMT-6 mammary tumor cells are more resistant to alkylating agents (cyclophosphamide, cisplatin, thiotepa) when grown in 3D spheroids (Kobayashi et al., 1993; Tan et al., 2006). Single or clustered cells detach from primary breast or prostate tumors and are then found in the bloodstream (Aceto et al., 2014). In the peritoneal, pleural or pericardial cavity, the formation and properties of these tumoral 3D aggregates, which occur in breast, ovarian, gastrointestinal, pancreatic and lung cancers, remain poorly understood. Their specific mechanisms of resistance to treatment are largely understudied. However, in peritoneal carcinomatosis of ovarian cancer patients, it is largely accepted that the peritoneal microenvironment contributes to chemotherapy resistance (Castells et al., 2012), in particular via mesenchymal stem cells (MSCs) (Roodhart et al., 2011). They have been shown to contribute to tumor cells’ ability to form spheroids called tumorospheres under anchorage-independent culture conditions, increasing their capacity to induce tumor formation in vivo after mouse xenografts (Kabashima-Niibe et al., 2013; Lee et al., 2018).

The class I PI3K (phosphoinositide 3-kinase)/Akt axis is one of the most commonly deregulated pathways in cancers, including in ovarian, breast and lung cancers. There are four heterodimeric isoforms of class I PI3K in vertebrates: α, β, γ and δ. PI3K activation results in the phosphorylation of phosphatidylinositol-4,5-biphosphate (PIP2) into phosphatidylinositol-3,4,5-triphosphate (PIP3), which acts as a second messenger in the cell. Downstream signals of PI3K, including the members of the Akt/mTOR (mammalian target of rapamycin) pathway, are numerous and involved in many cellular processes, including cell-cycle progression and actin cytoskeleton rearrangement (Baer et al., 2014; Tsujita and Itoh, 2015; Vanhaesebroeck et al., 2010; Zhang et al., 2017).

One PI3Kα-selective inhibitor, BYL719 (Alpelisib), was just approved for use in breast cancer with *PIK3CA* mutation (André et al., 2019; Mayer et al., 2017), and BYL719 in combination with estrogen receptor inhibitors show lower toxicity compared to the use of pan-PI3K inhibitors. PI3Kα/Akt/mTOR signaling pathway alterations and more specifically alterations in the frequency of *PIK3CA* mutations (*PIK3CA* encodes for the PI3Kα catalytic subunit) varied in non-small cell lung cancer (NSCLC) in published studies (Arcucci et al., 2021; Wu et al., 2016). In ovarian cancer, the PI3K signaling pathway is described as a predictor of the invasive and migratory potential of ovarian tumor cells (Bai et al., 2015). Mutations are found in the *PIK3CA* gene in 6%–33% of patients, and amplification or copy gain of this gene is found in 13%–39.8% of cases (Levine, 2005); these alterations are non-exclusive. Regarding the distributions of these alterations in different ovarian carcinoma subtypes, *PIK3CA* mutations are less common in high-grade serous carcinomas, while amplifications are frequently observed. The genetic landscape of ovarian carcinomas suggests that PI3K inhibitors, and in particular those targeting PI3Kα, could be tested in this pathology.

Twenty years ago, peritoneal carcinomatosis was seen as an incurable pathology requiring palliative care only. However, the introduction of loco-regional therapies combining cytoreductive surgery with hyperthermic intraperitoneal chemotherapy (HIPEC) based on platinum salts or taxanes changed the management of this disease (Wright et al., 2015). Signal-targeted therapies, such as those targeting the PI3K/Akt pathway, are currently being tested with the goal of improving disease-free survival rates (Mabuchi et al., 2015).

The importance of PI3Kα activity in tumoral aggregates from adenocarcinomas disseminated in serous cavity is unknown. Our objective is to demonstrate the involvement of PI3Kα during tumorosphere formation and maintenance in serous cavities and their role in chemotherapy resistance.

## Results

### Peritoneal tumor cells from ascites punctures form adhesive aggregates whose growth is delayed by a high number of initial cells

Ascites of 12 ovarian cancer patients were punctured and analyzed to detect the presence of adenocarcinomatous cells. We first observed tumor cell aggregates found in ascites using cytospin or an embedded preparation analyzed with standard immunohistochemistry techniques, Papanicolaou (PAP), May-Grünwald-Giemsa (MGG), periodic acid–Schiff (PAS) and epithelial cell adhesion molecule staining (Ber-EP4) (Fig. 1A). We observed heterogeneity in terms of the distribution of cellular aggregate density (Suppl. Table S1). Most ascites (11/12 patients) were composed of limited numbers of isolated tumoral cells and numerous aggregated tumoral cells (representative image, Fig. 1A left). Only one case revealed numerous isolated cells alongside aggregates (representative image, Fig. 1A right). When grown in 3D-favorable conditions, ex-vivo cultures of ascites cells formed dense and cohesive spheroids (Fig. 1B). Interestingly, ovarian cell lines derived from ascites were enriched in *PIK3CA* mutations and/or p53 alterations compared to primary tumor cell lines (Suppl. Table S2). To reproduce the carcinomatosis development of cohesive aggregates (Fig. 1C), we first developed a fluorescent 3D culture system of peritoneal tumor cells with *PIK3CA*-mutant ovarian cancer cell line SKOV-3 stably expressing GFP (Fig. 1D). After 14 days, we analyzed the influence of cell numbers in aggregates and observed an increase of 75% and 13% in the tumorosphere area when 5,000 and 10,000 cells were seeded, respectively (Fig. 1D, E). A higher number of initial cells appears to slow tumorosphere growth. Interestingly, area evolution (+24% in 7 days) (Fig. 1D, E) was associated with the constant expression of fluorescent GFP as a readout of cellular activity (Fig. 1F). As expected, fluorescence tends to decrease in the core of tumorospheres (Fig. 1D). Ascites is a mix of different cell populations (Fig. 1A). To reproduce this heterogeneity, we added a stromal compartment, adding MSC, known to be recruited at inflammatory tumoral sites. We tested their capacity to aggregate with tumor cells. We carried out 3D cultures with 10,000 MSC alone and observed a completely different aggregate evolution profile compared to SKOV-3 cells; there was very fast cell cohesion the day after seeding, and within 14 days, the aggregate area had decreased by more than 60% compared to the initial area (Suppl. Fig. S1A and B). We also observed the formation of fibrous structures within MSC aggregates, the nature of which remains to be determined (Suppl. Fig S1C). Those structures were also observed in aggregates developed from patient-derived tumor cells (Fig. 1B, top right). We then co-cultured 5000 GFP-expressing SKOV-3 cells with different quantities of Qdot-labelled MSCs (Fig. 1G-H) (Latifi et al., 2012). The heterotypic tumorosphere area decreased proportionally to the number of MSCs, with an apparent increased cohesion (Fig. 1G, H). GFP fluorescence corresponding to tumor cells did not vary regardless of MSC number, indicating that these cells induced stronger tumorosphere cohesion without modifying tumor cell proliferation (Fig. 1H, I). At a high ratio of MSCs to tumor cells, MSCs clustered in the center of tumorospheres and colocalized with tumor cells (yellow staining, Suppl. Fig. S1D).

**Figure 1:**
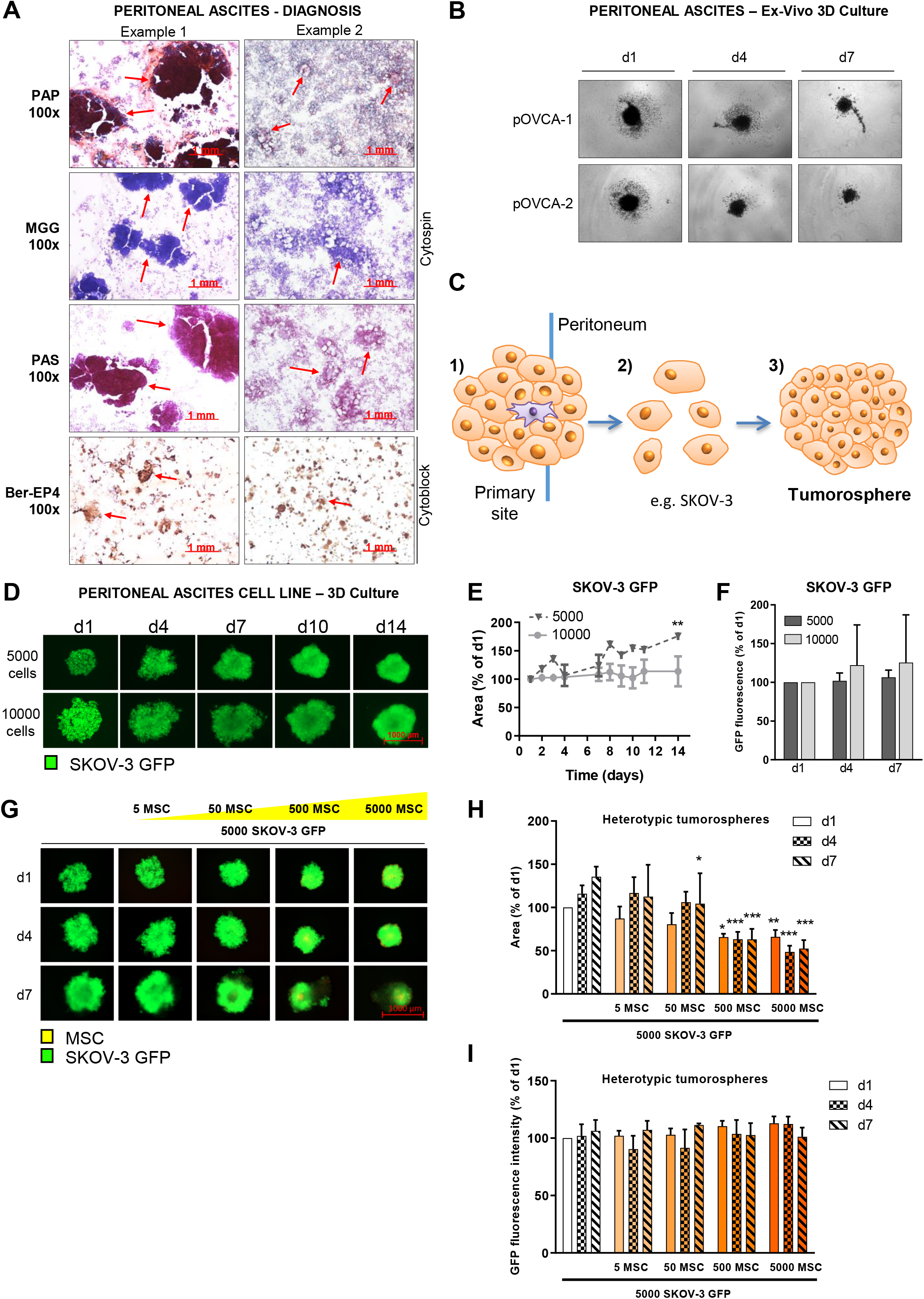
Modeling tumor aggregates derived from ovarian cancer ascites. (A) Immunohistochemistry of high-grade ovarian serous carcinoma from ascites fluid, in which tumorospheres are indicated by arrows (100x magnification). PAP = Papanicolaou, MGG = May-Grünwald-Giemsa, PAS = Periodic acid–Schiff. (B) Evolution of tumorosphere morphology 1, 4 or 7 days after seeding. (C) Model of dissemination in serous cavities from the primary tumor facilitated by the flow of liquid and formation of tumorospheres. (D) Morphology and (E) area of SKOV-3-GFP (green fluorescent protein) tumorospheres. Measurements with ZEN^®^ software (x5 magnification, fluorescence, n ≥ 2). Comparison of the tumorosphere area with 5000 or 10000 SKOV-3-GFP cells on day 14 by Student t-test p <0.01 (**). (F) GFP fluorescence intensity was evaluated at D1, D4 and D7 using ZEN^®^ software (in fluorescence, x5 magnification, n = 3). (G) Morphology of heterotypic tumorospheres comprising 5000 SKOV-3-GFP cells (green) cultured with or without different numbers of mesenchymal stem cells (MSCs) (5, 50, 500, 5000) 1, 4 or 7 days after seeding. (H) Area and (I) GFP fluorescence intensity were evaluated at D1, D4 and D7 using ZEN^®^ software (in fluorescence, x5 magnification, n = 3). Mean +/− SEM (ANOVA test comparison with the condition without MSCs at the corresponding observation day, p < 0.001 (***), p < 0.01 (**), p < 0.05 (*)).

These results reveal that small-sized tumorospheres exhibit cellular growth and that the addition of MSCs to tumor cell spheroids results in increased cohesion of tumorospheres, reminiscent of the observed formation of tumorospheres from patients’ ascites cells.

### PI3Kα inhibition prevents cohesion of ovarian cancer cell tumorospheres

The PI3K/Akt pathway plays a role in cell proliferation and survival but is also involved in actin cytoskeleton regulation, suggesting a role in tumorosphere formation (Baer et al., 2014; Tsujita and Itoh, 2015; Vanhaesebroeck et al., 2010). We confirmed a concentration-dependent inhibition of Akt phosphorylation on S473 and T308 after A66 and BYL719 treatment (Fig. 2A-C), indicating 1) that the PI3K pathway is constitutively activated in SKOV-3 cells, 2) that this activation is highly dependent on the PI3Kα isoform and 3) that the pharmacological agents used prevent its activity.

**Figure 2:**
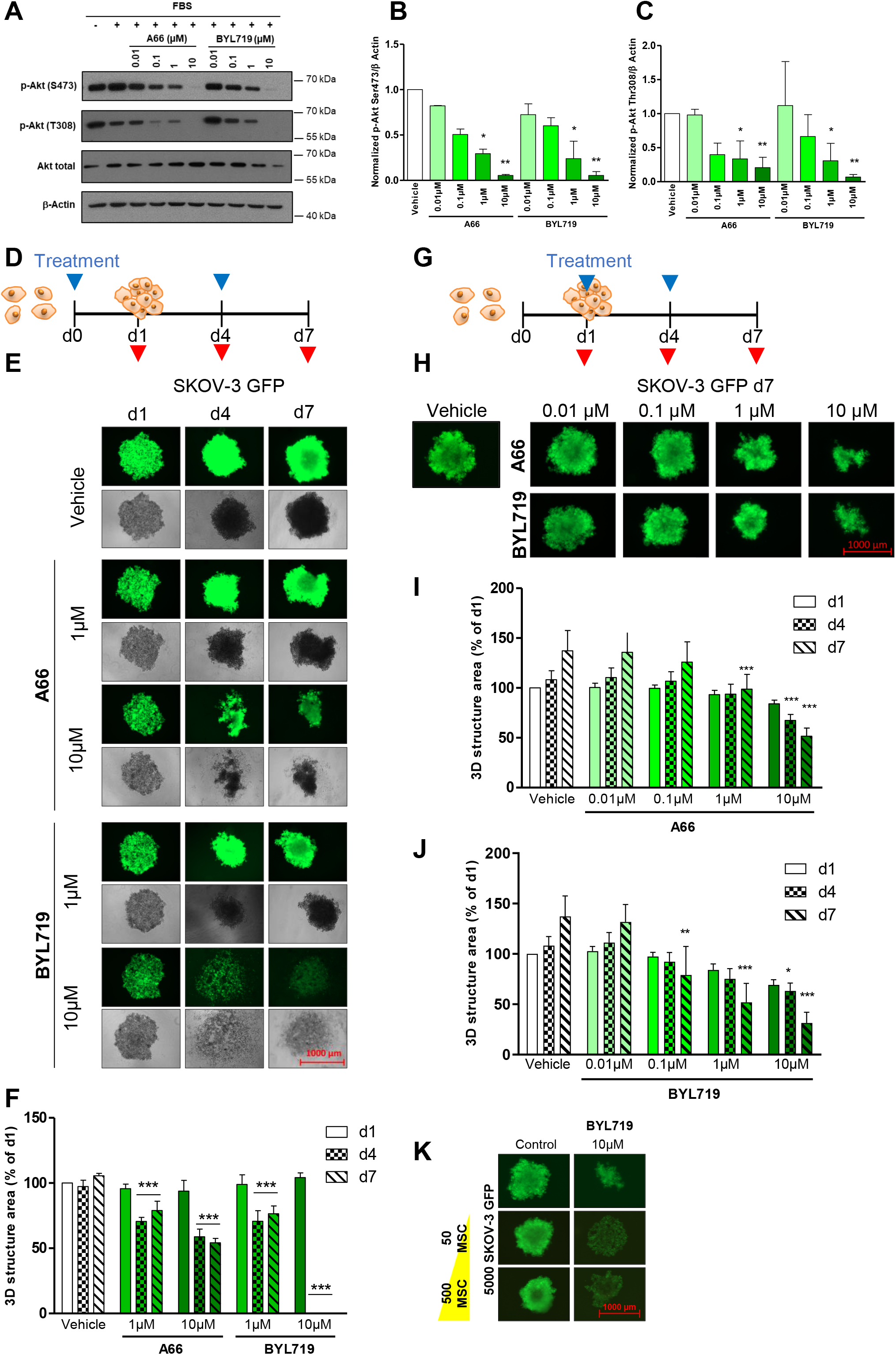
PI3Kα inhibition prevents formation and maintenance of ovarian cancer cell tumorospheres. (A) SKOV-3 cells were serum-deprived overnight, pre-treated for 1 h with specific PI3Kα inhibitors and then serum-activated for 15 minutes. Actin-standardized quantification of the amount of (B) phospho-Akt (S473) and (C) phospho-Akt (T308) relative to the vehicle condition (without treatment). (D) 5,000 SKOV-3-GFP cells that will form tumoropheres at D1 were treated or not at D0 and D4 with A66 and BYL719 (1 and 10 μM). (E, F) SKOV-3-GFP tumorospheres were treated according to the schedule in (D) and morphology was observed 1, 4 or 7 days after seeding. (F) Area was evaluated at D1, D4 and D7 using ZEN^®^ software (x5 magnification), n ≥ 3, mean +/− SEM (ANOVA test comparison with the vehicle condition of the corresponding observation day, p <0.001 (***), p <0.01 (**), p <0.05 (*)). (G) Tumorospheres formed of 5,000 SKOV-3-GFP cells were treated or not at D1 and D4 with A66 or BYL719 (0.01, 0.1, 1 and 10 μM). (H) Tumorosphere morphology was evaluated on D1, D4 and D7. (I, J) Tumorosphere area was evaluated on D1, D4 and D7 using ZEN^®^ software (x5 magnification), n ≥ 3, mean +/− SEM (ANOVA test comparison with the control condition of the corresponding observation day, p <0.001 (***), p <0.01 (**), p <0.05 (*)). (K) Tumorospheres formed of 5,000 SKOV-3-GFP cells (green) associated with different ratios of MSCs (labeled in red using Quantum Dots) were produced and treated with 10 μM BYL719. Tumorosphere morphology was evaluated using ZEN^®^ software (in fluorescence, x5 magnification) on D7.

To determine the action of this pathway in the formation of SKOV-3-GFP multicellular aggregates, we treated them at seeding and 3 days later with the PI3Kα-selective inhibitors A66 and BYL719 (Fig. 2D). We next compared the efficiency of a low (1 μM) and medium (10 μM) dose of PI3K inhibitors and observed the morphology of the 3D multicellular structures. After 4 and 7 days of treatment with 10 μM of BYL719, we observed a complete reduction of the tumorosphere area (Fig. 2E–F). Aggregates treated with 10 μM of A66 or 1 μM of BYL719 had comparable evolution profiles (Fig. 2E), with a decrease in tumorosphere area of about 30% compared to the control at day 4 (Fig. 2F). This effect was maintained over time.

To measure PI3Kα inhibitor effects on tumorosphere maintenance, we treated already established SKOV-3-GFP cell aggregates (Fig. 2G). PI3Kα inhibitors caused a concentration-dependent decrease in tumorosphere area and a tumorosphere morphology change with maximum effect at day 7 (Fig. 2H-J). Used at 1 μM, treatment with BYL719 was more effective than A66; there was a significant decrease in tumorosphere area of 86% and 43%, respectively, compared to the vehicle at day 7 (Fig. 2I and J). This corresponded to a decrease in the initial tumorosphere size.

Finally, adding MSCs, which compacted the tumorospheres, to BYL-719 at 10 μM reduced tumorosphere cohesion (Fig. 2K). The absence of a direct effect of all PI3K inhibitors on MSCs was verified (Suppl. Fig. S1E, F).

PI3Kα inhibitors prevent cohesive tumorosphere formation and maintenance in an α isoform-specific manner.

### PI3Kα inhibition impacts multicellular tumor cell viability in 3D cultures

To extend our findings, we used 2 other cell lines derived from serous cavities presenting *PIK3CA* mutations: the lung cancer cell line H460 and the breast cancer cell line MCF-7. Using a survival MTT assay, we first verified that they were sensitive in 2D to PI3Kα inhibitors (Fig. 3A). Heterogeneous responses to PI3K inhibitors were observed (Fig. 3A); the H460 cell line appeared to be resistant to A66 and to low concentrations of BYL719 (Fig. 3A, middle).

**Figure 3:**
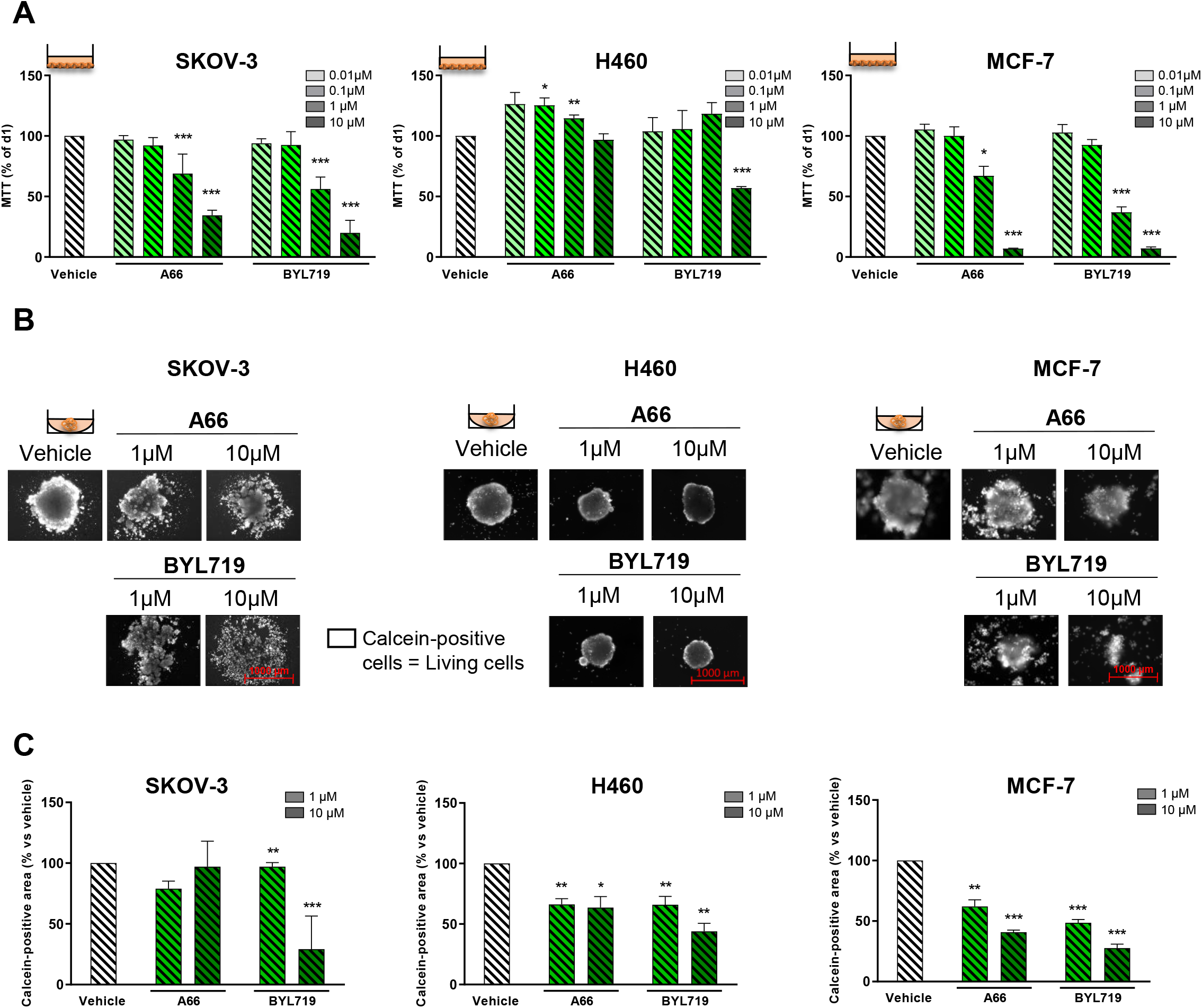
PI3Kα inhibition impacts multicellular cell viability. (A) SKOV-3, H460 and MCF-7 cell monolayers were treated or not with A66 or BYL791 and living cells were quantified after 3 days with a MTT colorimetric assay. (B-C) Tumorospheres formed of 5,000 SKOV-3-GFP, H460 and MCF-7 cells were treated or not at D1 and D4 with A66 or BYL719. (B) Tumorosphere morphology was evaluated on D7 using ZEN^®^ software (in fluorescence, x5 magnification); living cells were labeled with calcein (white), n = 3. (C) The area of living cells after treatment was quantified using ImageJ^®^ software, n = 3. Mean +/− SEM (ANOVA test comparison with the control condition of the corresponding observation day, p <0.001 (***), p <0.01 (**), p <0.05 (*)).

We next aimed to determine the impact of the 3D culture on PI3Kα. Inhibitor efficiency. In particular, we were interested in analyzing the organization and distribution of living and dead cells within tumorospheres under treatment. PI3K inhibitor treatment altered tumorosphere cohesion in SKOV-3 and MCF-7 cells (Fig. 3B, Suppl. Fig. S2A). We quantified the living cell areas using calcein staining and demonstrated that BYL719 significantly decreased this marker in all tested cell lines (Fig. 3B-C, Suppl. Fig S2A). SKOV-3 and H460 cell treatment with PI3Kα inhibitors tended to increase the number of dead cells (Suppl. Fig. S2B). Just 10 μM BYL719 caused a significant decrease in the living cell area of all cell lines (Fig. 3B, C). PI3Kα inhibitors induced a rapid decrease in tumorosphere area and viability.

### PI3K inhibitors decrease tumorosphere survival in the presence of cisplatin

If used in the clinic to treat ovarian cancer, PI3K inhibitors will be tested in combination (Pons-Tostivint et al., 2017). A combination of a low concentration (0.05 or 0.5 μM) of cisplatin with PI3Kα inhibitors appeared to alter more efficiently the SKOV-3-GFP tumorosphere area and living cells compared to the control (cisplatin alone), but not significantly more than the corresponding inhibitor alone (Fig. 4A–D). At low doses of cisplatin, both PI3K inhibitors significantly decreased the 3D structure area (Fig. 4C). The addition of 1 μM PI3Kα inhibitors (A66 or BYL719) to a high concentration of cisplatin (5 μM) triggered a huge decrease in living cells compared to each agent alone, suggesting an additive effect for both drugs (Fig. 4D). Differences in the effects observed with these different assays show that the key combined action of PI3Kα inactivation and cisplatin treatment consists in decreasing cell survival and tumorosphere area.

**Figure 4:**
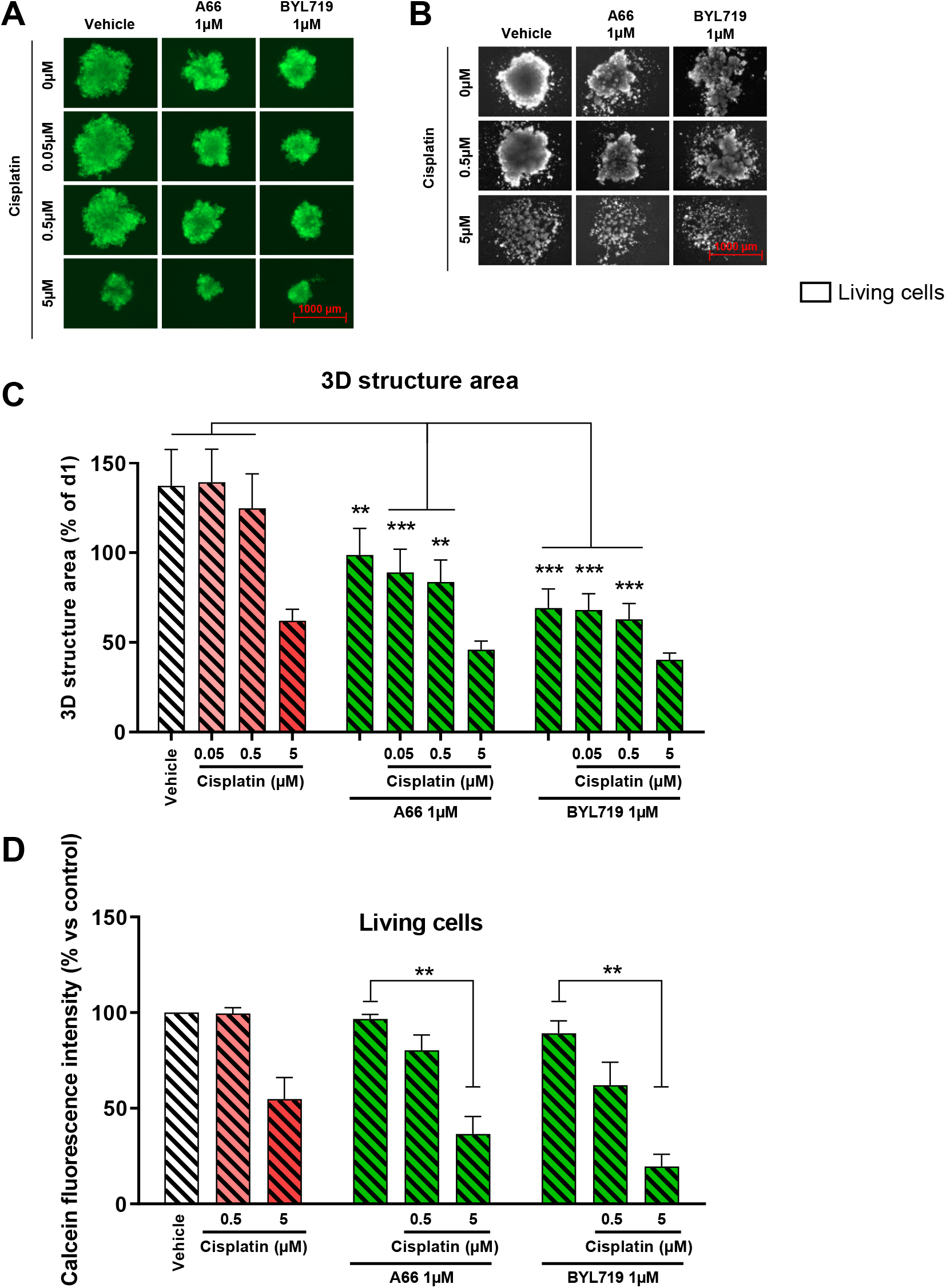
PI3K inhibitors decrease tumorosphere survival in the presence of cisplatin. Tumorospheres formed of 5,000 SKOV-3 cells were treated or not on D1 and D4 after seeding with a fixed concentration of 1 μM A66, BYL719, TGX-221 or AZD8186, in combination or not with increasing concentrations of cisplatin (0.05, 0.5, 5 μM). (A) Tumorosphere morphology and (C) area (n ≥ 3) were evaluated using ZEN^®^ software (in fluorescence, x5 magnification). (B) Living cells were marked with calcein and (D) the living cell intensity after treatment was quantified using ImageJ^®^ software (n = 2), mean +/− SEM (ANOVA test comparison with cisplatin concentration and the control of the corresponding observation day, p <0.001 (***), p <0.01 (**), p <0.05 (*)).

Compared to cisplatin treatment, treatment with PI3Kα inhibitors results in decreased tumorosphere survival.

## Discussion

Solid cancers associated with peritoneal, pleural or pericardial carcinomatosis frequently develop fluid accumulations called ascites in the case of the peritoneum. This early step in the metastatic process involves tumor cell aggregation within the peritoneal cavity. This 3D structure enhances the ability of tumor cells to survive conventional chemotherapies (Liao et al., 2014; Tan et al., 2006), and we found that PI3Kα activation through its oncogenic mutations contributes to this process.

PI3Kα inhibitors are promising therapeutic tools to prevent local dissemination in serous cavities. The activating mutations of *PIK3CA* and the disabling mutation of *PTEN* have been identified with p53 mutation as the three most frequent point mutations among 21 types of cancers. Several studies using genetically engineered mouse models have demonstrated the role of mutant *PIK3CA* in tumor initiation, progression and maintenance (Arcucci et al., 2021). Despite these data, given the mild effect of PI3Kα inhibitors as clinical monotherapy, the oncogenic potential of mutated PI3Kα is challenged. Analyzing the action of mutant PI3Kα in combination with other genetic alterations in patients and preclinical models is of utmost importance to better understand the cooperation mechanisms between different oncogenic processes and provide novel therapeutic solutions (Pons-Tostivint et al., 2017). p53 mutations are also known to increase *PIK3CA* promoter activity (Gaikwad et al., 2013), contributing to global PI3K/Akt pathway activation in cell lines coming from ascites. Globally, it seems that compared to primary tumors, cell lines coming from ascites are enriched in either mutant PIK3CA and/or genetic p53 inactivation (Table S2).

PI3K, and more particularly the α isoform, plays a role in actin cytoskeleton rearrangement (Baer et al., 2014; Campa et al., 2015). Actin involvement in supporting adhesive connections is well established and, according to our hypothesis, PI3Kα activity would facilitate ovarian tumor cell aggregate formation (Tsujita and Itoh, 2015; Vanhaesebroeck et al., 2010).

To validate this hypothesis, we treated SKOV-3 cell tumorospheres during seeding with different concentrations of the PI3Kα. inhibitors A66 (Jamieson et al., 2011) and BYL719 (Fritsch et al., 2014). After treatment of these aggregates, microscopy revealed sphere disorganization. Complete disintegration was seen following treatment with 10 μM BYL719. PI3Kα seems to play a role in both ovarian tumor cell aggregate formation and maintenance.

PI3Kα action on the actin cytoskeleton could be the source of our demonstrated effect of PI3K inhibitors on spheroids (Lien et al., 2017). This hypothesis is reinforced by the knowledge of an existing link between polymerized actin and adhesive molecules like E- or P-cadherins. P-cadherin expression promotes the aggregation of circulating ovarian cancer cells (Usui et al., 2014). These elements highlighting the role of actin could in part explain PI3Kα involvement in cell clustering (van Baal et al., 2018; Saias et al., 2015). Therapies targeting adhesion between cancer cells of the same type, such as PI3Kα inhibitors, are worth consideration for improving peritoneal carcinomatosis management.

It has been repeatedly demonstrated that 2D cytotoxicity studies do not account for complete tissue physiology and overestimate the cytotoxic potential of tested molecules (Kobayashi et al., 1993; Shield et al., 2009). In a 3D culture, these cells are 18, 15 or 58 times more resistant to cisplatin at 25, 50 and 100 μM, respectively (Kobayashi et al., 1993). In addition, proliferation within the spheres decreases in comparison with cells cultured in 2D, partially explaining why these cells can survive therapies that induce death in replicating cells (Burleson et al., 2004; Kobayashi et al., 1993). Here, all cells cultured in 3D remain sensitive to PI3K inhibition, suggesting that PI3K inhibition impacts cells with reduced proliferation.

Tumor aggregate disorganization induced by PI3Kα inhibitors could allow better cisplatin bioavailability within tumorospheres by breaking the decreasing diffusion gradient from the periphery to the center of these 3D structures (Sant and Johnston, 2017). Tumorosphere dissociation could also decrease the solid stress that builds up in 3D culture (Delarue et al., 2014), which was shown to decrease cell proliferation and chemotherapy efficiency (Rizzuti et al., 2020). Other parameters we could have analyzed are the response to increased liquid pressure (Klymenko et al., 2018) or to shear stress caused by ascites fluid motion in the peritoneal cavity (Rizvi et al., 2013). They were shown by others to increase tumor cell proliferation rate and they could allow cisplatin to function.

We tested the role of oncogenic PI3Kα in different types of adenocarcinomas with dissemination in serous fluid, such as ovary, lung and breast cancers. Our findings are potentially of therapeutic importance in ovarian cancer, which at diagnosis most often presents an advanced stage associated with peritoneal dissemination. The current therapeutic arsenal lacks treatments targeting the more advanced disease, which represents a major public health issue due to its morbidity and mortality. To date, HIPEC is of therapeutic interest for the treatment of peritoneal carcinomatosis, but its use remains infrequent because of potential toxicities (van Driel et al., 2018; Goéré et al., 2013). Our data are in favor of adding PI3Kα inhibitors to this therapeutic arsenal.

## Acknowledgements

We thank all members of SigDYN for their comments during the writing of the manuscript, the CRCT core technology platform, particularly Laetitia Ligat, for imaging and Marie Veronique Joubert for her help in MSC culture, the Gynecology Oncology department in IUCT-O for providing patient ascites. JGG’s laboratory is part of TOUCAN (Laboratoire d’Excellence, ANR), an integrated research program on signal-targeted drug resistance. For this topic, JGG’s laboratory received funding from the ARC (Foundation for Cancer Research) (PJA20171206596), TOUCAN (Laboratoire d’Excellence, ANR), Fondation de France (BT’s salary) and Hôpitaux de Toulouse (AT’s salary).

## Author contributions

**Benoît Thibault:** conceptualization; data curation; formal analysis; investigation; methodology; validation; visualization; writing – review & editing. **Adrien Thole:** data curation; formal analysis; investigation; methodology; visualization; writing – original draft; review & editing. **Céline Basset:** data curation; formal analysis; methodology; resources; visualization; writing – review & editing. **Julie Guillermet-Guibert:** conceptualization; data curation; formal analysis; funding acquisition; investigation; methodology; project administration; supervision; validation; visualization; writing – original draft; writing – review & editing.

## MATERIAL AND METHODS

### Patient ascites-derived cell tumorospheres

Ascites fluids were taken from 12 patients with ovarian cancer after consent (CRB Cancer des Hôpitaux de Toulouse, IUCT-O, Toulouse – BB-0033-00014, DC-2008-463, AC-2013-1955). Supplementary Table S1 details patient information; clinical and biological annotations of the samples have been declared to the CNIL (Comité National Informatique et Libertés).

pOVCA-1 and pOVCA-2 were ovarian tumor cells isolated from patients’ ascites and cultivated in DMEM supplemented with 10% FCS (fetal calf serum), 1% L-glutamine, 1% penicillin-streptomycin and 0.01% plasmocin (Invitrogen).

### Cell lines, 3D culture, treatments and phenotypic analysis

SKOV-3-GFP cells correspond to genetically modified SKOV-3 wild-type (WT) cells expressing GFP. MSCs were derived from healthy donors who underwent hip surgery (Etablissement Français du Sang). Cells were cultured at 37°C, 5% CO_2_ in a humid atmosphere, in RPMI 1640 medium (Sigma) for SKOV-3 and H460 cells, or DMEM (Sigma) for MSCs and MCF-7 cells; 50% of each medium was used when MSCs and SKOV-3 cells were mixed. The culture medium was supplemented with 10% FCS (fetal calf serum), 1% L-glutamine, 1% penicillin-streptomycin and 0.01% plasmocin (Invitrogen).

One thousand patient-derived tumoral cells or 5000 SKOV-3-GFP, SKOV-3 WT cells, H460 or MCF-7 cells with or without increasing numbers of MSCs were seeded in round-bottom 96-well plates (Nunclon Sphera, ThermoFisher Scientific). Living cell numbers and metabolic activity were measured with an MTT assay or labeled for 1 h using the LIVE/DEAD^®^ kit (ThermoFisher Scientific) containing 2 μM calcein and 4 μM EthD-1 (Ethidium homodimer-1). The cell clusters obtained were observed and imaged with an AxioVert microscope on days 1, 4, 7, 10 and 14 using ZEN^®^ software (Carl Zeiss). The same software measured their diameter and surface. For each condition, triplicates were performed and the average area was compared to the day 1 average of the corresponding experiment (representing 100% of the area). Western blotting was performed as described in our previous work (Baer et al., 2014) with the following antibodies:

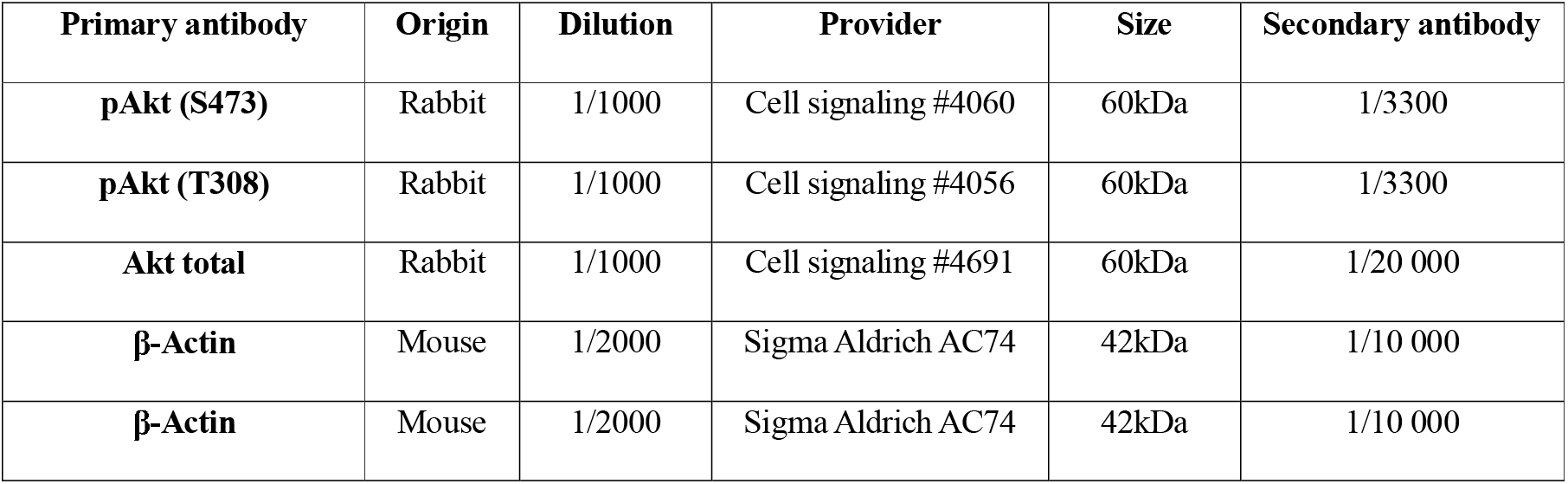

| Primary antibody | Origin | Dilution | Provider | Size | Secondary antibody |
| --- | --- | --- | --- | --- | --- |
| **pAkt (S473)** | Rabbit | 1/1000 | Cell signaling #4060 | 60kDa | 1/3300 |
| **pAkt (T308)** | Rabbit | 1/1000 | Cell signaling #4056 | 60kDa | 1/3300 |
| **Akt total** | Rabbit | 1/1000 | Cell signaling #4691 | 60kDa | 1/20 000 |
| **β-Actin** | Mouse | 1/2000 | Sigma Aldrich AC74 | 42kDa | 1/10 000 |
| **β-Actin** | Mouse | 1/2000 | Sigma Aldrich AC74 | 42kDa | 1/10 000 |

### Pharmacological inhibitors

Treatments used in this study were as follows: Cisplatin (kindly supplied by Institut Universitaire du Cancer – Oncopole (IUCT-O)), A66 (Axon Medchem) PI3Kα inhibitor (in vitro IC50 in nM: p110α: 32; β: >12500; δ: >1250; γ: 3480) (Jamieson et al., 2011); BYL719 (Apex Bio) PI3Kα inhibitor (in vitro IC50 in nM: p110α 4.6; β: 1156; δ: 290; γ: 250) (Fritsch et al., 2014).

### Qdot^®^ cell labeling of MSCs

Labeling was performed according to the supplier’s recommendations, with the equimolar mix from the labeling kit at dilution 1/100 incubated for 1 h at room temperature applied directly to the cell cultures for 45 minutes.

### Statistics

Quantitative variables are presented by means and qualitative data by percentages. Group comparisons were made using the ANOVA test (matched data) and Student’s t-test for parametric tests. For all experiments, a statistical difference is reached for p<0.05.

---

**Supplementary Figure S1: Development of heterotypic tumorospheres**. (A) Area and (B) morphology of MSC tumorospheres. Mean +/− SEM measurements with ZEN^®^ software (x5 magnification, n ≥ 2). (C) Observation of the formation of fibrous structures within MSC aggregates. (D) Observation of heterotypic tumorospheres of SKOV-3 cells (green) and MSCs (red). 5000 SKOV-3-GFP cells associated with different MSC ratios (500 or 5000) were observed at D1 and D7. (E, F) Spheres formed of 5000 MSCs were produced and treated or not with A66, BYL719 or cisplatin 24 h after seeding. (E) Morphology and (F) area of MSC tumorospheres were measured using ZEN^®^ software (x5 magnification, n = 3).

**Supplementary Figure S2: Effects of PI3K inhibitors on SKOV-3, H460 and MCF-7 tumorospheres.** Tumorospheres formed of 5000 SKOV-3, H460 and MCF-7 cells were produced and treated or not with A66, BYL719 at D1 and D4 after seeding. (A) At D7, living cells were labeled with calcein (green) and dead cells with EthD-1 (red) using the LIVE/DEAD^®^ kit. (B) The ratio of dead cells’ fluorescence on living cells’ fluorescence and compared to the vehicle condition. Mean +/− SEM, n = 3 (Student’s t-test, comparison with the vehicle condition, p <0.01 (**)).

**Supplementary Table S1: Patient data.** Data corresponding to the ascites samples isolated from patients. Patient number, age (in years), histology, aggregate numbers (per microscopy field) are represented.

**Supplementary Table S2: Frequency of *PIK3CA* or p53 mutations in cell lines derived from primary tumors versus serous fluid cavities, example of ovarian carcinoma. (A)** PIK3CA and p53 mutational status in ovarian cancer cell lines coming from a primary tumor (tumor) or ascites fluid (ascites). The exact mutation is indicated when known. WT = wild type. (B) Frequency of ovarian cancer cell lines harboring a PIK3CA and/or p53 mutation depending on their origin (primary tumor or ascites). Mainly identified from Cellosaurus or according to the literature (Beaufort et al., 2014).

## References

Aceto, N., Bardia, A., Miyamoto, D.T., Donaldson, M.C., Wittner, B.S., Spencer, J.A., Yu, M., Pely, A., Engstrom, A., Zhu, H., et al. (2014). Circulating tumor cell clusters are oligoclonal precursors of breast cancer metastasis. Cell 158, 1110–1122.

André, F., Ciruelos, E., Rubovszky, G., Campone, M., Loibl, S., Rugo, H.S., Iwata, H., Conte, P., Mayer, I.A., Kaufman, B., et al. (2019). Alpelisib for *PIK3CA*-Mutated, Hormone Receptor–Positive Advanced Breast Cancer. N. Engl. J. Med. 380, 1929–1940.

Arcucci, S., Ramos-Delgado, F., Cayron, C., Therville, N., Gratacap, M.-P., Basset, C., Thibault, B., and Guillermet-Guibert, J. (2021). Organismal roles for the PI3Kα and β isoforms: their specificity, redundancy or cooperation is context-dependent. Biochem. J. 478, 1199–1225.

van Baal, J.O.A.M., van Noorden, C.J.F., Nieuwland, R., Van de Vijver, K.K., Sturk, A., van Driel, W.J., Kenter, G.G., and Lok, C.A.R. (2018). Development of Peritoneal Carcinomatosis in Epithelial Ovarian Cancer: A Review. J. Histochem. Cytochem. 66, 67–83.

Baer, R., Cintas, C., Dufresne, M., Cassant-Sourdy, S., Schönhuber, N., Planque, L., Lulka, H., Couderc, B., Bousquet, C., Garmy-Susini, B., et al. (2014). Pancreatic cell plasticity and cancer initiation induced by oncogenic Kras is completely dependent on wild-type PI 3-kinase p110α. Genes Dev. 28, 2621–2635.

Bai, H., Li, H., Li, W., Gui, T., Yang, J., Cao, D., and Shen, K. (2015). The PI3K/AKT/mTOR pathway is a potential predictor of distinct invasive and migratory capacities in human ovarian cancer cell lines. Oncotarget 6.

Beaufort, C.M., Helmijr, J.C.A., Piskorz, A.M., Hoogstraat, M., Ruigrok-Ritstier, K., Besselink, N., Murtaza, M., van IJcken, W.F.J., Heine, A.A.J., Smid, M., et al. (2014). Ovarian cancer cell line panel (OCCP): clinical importance of in vitro morphological subtypes. PloS One 9, e103988.

Burleson, K.M., Casey, R.C., Skubitz, K.M., Pambuccian, S.E., Oegema, T.R., and Skubitz, A.P.N. (2004). Ovarian carcinoma ascites spheroids adhere to extracellular matrix components and mesothelial cell monolayers. Gynecol. Oncol. 93, 170–181.

Campa, C.C., Ciraolo, E., Ghigo, A., Germena, G., and Hirsch, E. (2015). Crossroads of PI3K and Rac pathways. Small GTPases 6, 71–80.

Castells, M., Thibault, B., Delord, J.-P., and Couderc, B. (2012). Implication of Tumor Microenvironment in Chemoresistance: Tumor-Associated Stromal Cells Protect Tumor Cells from Cell Death. Int. J. Mol. Sci. 13, 9545–9571.

Delarue, M., Montel, F., Vignjevic, D., Prost, J., Joanny, J.-F., and Cappello, G. (2014). Compressive Stress Inhibits Proliferation in Tumor Spheroids through a Volume Limitation. Biophys. J. 107, 1821–1828.

van Driel, W.J., Koole, S.N., Sikorska, K., Schagen van Leeuwen, J.H., Schreuder, H.W.R., Hermans, R.H.M., de Hingh, I.H.J.T., van der Velden, J., Arts, H.J., Massuger, L.F.A.G., et al. (2018). Hyperthermic Intraperitoneal Chemotherapy in Ovarian Cancer. N. Engl. J. Med. 378, 230–240.

Fritsch, C., Huang, A., Chatenay-Rivauday, C., Schnell, C., Reddy, A., Liu, M., Kauffmann, A., Guthy, D., Erdmann, D., De Pover, A., et al. (2014). Characterization of the Novel and Specific PI3K Inhibitor NVP-BYL719 and Development of the Patient Stratification Strategy for Clinical Trials. Mol. Cancer Ther. 13, 1117–1129.

Gaikwad, S.M., Gunjal, L., Junutula, A.R., Astanehe, A., Gambhir, S.S., and Ray, P. (2013). Non-Invasive Imaging of Phosphoinositide-3-Kinase-Catalytic-Subunit-Alpha (PIK3CA) Promoter Modulation in Small Animal Models. PLoS ONE 8, e55971.

Goéré, D., Malka, D., Tzanis, D., Gava, V., Boige, V., Eveno, C., Maggiori, L., Dumont, F., Ducreux, M., and Elias, D. (2013). Is there a possibility of a cure in patients with colorectal peritoneal carcinomatosis amenable to complete cytoreductive surgery and intraperitoneal chemotherapy? Ann. Surg. 257, 1065–1071.

Jamieson, S., Flanagan, J.U., Kolekar, S., Buchanan, C., Kendall, J.D., Lee, W.-J., Rewcastle, G.W., Denny, W.A., Singh, R., Dickson, J., et al. (2011). A drug targeting only p110α can block phosphoinositide 3-kinase signalling and tumour growth in certain cell types. Biochem. J. 438, 53–62.

Kabashima-Niibe, A., Higuchi, H., Takaishi, H., Masugi, Y., Matsuzaki, Y., Mabuchi, Y., Funakoshi, S., Adachi, M., Hamamoto, Y., Kawachi, S., et al. (2013). Mesenchymal stem cells regulate epithelial-mesenchymal transition and tumor progression of pancreatic cancer cells. Cancer Sci. 104, 157–164.

Klymenko, Y., Wates, R.B., Weiss-Bilka, H., Lombard, R., Liu, Y., Campbell, L., Kim, O., Wagner, D., Ravosa, M.J., and Stack, M.S. (2018). Modeling the effect of ascites-induced compression on ovarian cancer multicellular aggregates. Dis. Model. Mech. 11, dmm034199.

Kobayashi, H., Man, S., Graham, C.H., Kapitain, S.J., Teicher, B.A., and Kerbel, R.S. (1993). Acquired multicellular-mediated resistance to alkylating agents in cancer. Proc. Natl. Acad. Sci. U. S. A. 90, 3294–3298.

Latifi, A., Luwor, R.B., Bilandzic, M., Nazaretian, S., Stenvers, K., Pyman, J., Zhu, H., Thompson, E. W., Quinn, M.A., Findlay, J.K., et al. (2012). Isolation and Characterization of Tumor Cells from the Ascites of Ovarian Cancer Patients: Molecular Phenotype of Chemoresistant Ovarian Tumors. PLoS ONE 7, e46858.

Lee, J.-H., Kim, S.-K., Khawar, I.A., Jeong, S.-Y., Chung, S., and Kuh, H.-J. (2018). Microfluidic co-culture of pancreatic tumor spheroids with stellate cells as a novel 3D model for investigation of stroma-mediated cell motility and drug resistance. J. Exp. Clin. Cancer Res. 37.

Levine, D.A. (2005). Frequent Mutation of the PIK3CA Gene in Ovarian and Breast Cancers. Clin. Cancer Res. 11, 2875–2878.

Liao, J., Qian, F., Tchabo, N., Mhawech-Fauceglia, P., Beck, A., Qian, Z., Wang, X., Huss, W.J., Lele, S.B., Morrison, C.D., et al. (2014). Ovarian Cancer Spheroid Cells with Stem Cell-Like Properties Contribute to Tumor Generation, Metastasis and Chemotherapy Resistance through Hypoxia-Resistant Metabolism. PLoS ONE 9, e84941.

Lien, E.C., Dibble, C.C., and Toker, A. (2017). PI3K signaling in cancer: beyond AKT. Curr. Opin. Cell Biol. 45, 62–71.

Mabuchi, S., Kuroda, H., Takahashi, R., and Sasano, T. (2015). The PI3K/AKT/mTOR pathway as a therapeutic target in ovarian cancer. Gynecol. Oncol. 137, 173–179.

Mayer, I.A., Abramson, V.G., Formisano, L., Balko, J.M., Estrada, M.V., Sanders, M.E., Juric, D., Solit, D., Berger, M.F., Won, H.H., et al. (2017). A Phase Ib Study of Alpelisib (BYL719), a PI3Kα-Specific Inhibitor, with Letrozole in ER + /HER2 - Metastatic Breast Cancer. Clin. Cancer Res. 23, 26–34.

Pons-Tostivint, E., Thibault, B., and Guillermet-Guibert, J. (2017). Targeting PI3K Signaling in Combination Cancer Therapy. Trends Cancer 3, 454–469.

Rizvi, I., Gurkan, U.A., Tasoglu, S., Alagic, N., Celli, J.P., Mensah, L.B., Mai, Z., Demirci, U., and Hasan, T. (2013). Flow induces epithelial-mesenchymal transition, cellular heterogeneity and biomarker modulation in 3D ovarian cancer nodules. Proc. Natl. Acad. Sci. 110, E1974–E1983.

Rizzuti, I.F., Mascheroni, P., Arcucci, S., Ben-Mériem, Z., Prunet, A., Barentin, C., Rivière, C., Delanoë-Ayari, H., Hatzikirou, H., Guillermet-Guibert, J., et al. (2020). Mechanical Control of Cell Proliferation Increases Resistance to Chemotherapeutic Agents. Phys. Rev. Lett. 125, 128103.

Roodhart, J.M.L., Daenen, L.G.M., Stigter, E.C.A., Prins, H.-J., Gerrits, J., Houthuijzen, J.M., Gerritsen, M.G., Schipper, H.S., Backer, M.J.G., van Amersfoort, M., et al. (2011). Mesenchymal Stem Cells Induce Resistance to Chemotherapy through the Release of Platinum-Induced Fatty Acids. Cancer Cell 20, 370–383.

Saias, L., Gomes, A., Cazales, M., Ducommun, B., and Lobjois, V. (2015). Cell-Cell Adhesion and Cytoskeleton Tension Oppose Each Other in Regulating Tumor Cell Aggregation. Cancer Res. 75, 2426–2433.

Sant, S., and Johnston, P.A. (2017). The production of 3D tumor spheroids for cancer drug discovery. Drug Discov. Today Technol. 23, 27–36.

Shield, K., Ackland, M.L., Ahmed, N., and Rice, G.E. (2009). Multicellular spheroids in ovarian cancer metastases: Biology and pathology. Gynecol. Oncol. 113, 143–148.

Tan, D.S.P., Agarwal, R., and Kaye, S.B. (2006). Mechanisms of transcoelomic metastasis in ovarian cancer. Lancet Oncol. 7, 925–934.

Tsujita, K., and Itoh, T. (2015). Phosphoinositides in the regulation of actin cortex and cell migration. Biochim. Biophys. Acta 1851, 824–831.

Usui, A., Ko, S.Y., Barengo, N., and Naora, H. (2014). P-Cadherin Promotes Ovarian Cancer Dissemination Through Tumor Cell Aggregation and Tumor–Peritoneum Interactions. Mol. Cancer Res. 12, 504–513.

Vanhaesebroeck, B., Guillermet-Guibert, J., Graupera, M., and Bilanges, B. (2010). The emerging mechanisms of isoform-specific PI3K signalling. Nat. Rev. Mol. Cell Biol. 11, 329–341.

Wright, A.A., Cronin, A., Milne, D.E., Bookman, M.A., Burger, R.A., Cohn, D.E., Cristea, M.C., Griggs, J.J., Keating, N.L., Levenback, C.F., et al. (2015). Use and Effectiveness of Intraperitoneal Chemotherapy for Treatment of Ovarian Cancer. J. Clin. Oncol. 33, 2841–2847.

Wu, S.-G., Chang, Y.-L., Yu, C.-J., Yang, P.-C., and Shih, J.-Y. (2016). The Role of PIK3CA Mutations among Lung Adenocarcinoma Patients with Primary and Acquired Resistance to EGFR Tyrosine Kinase Inhibition. Sci. Rep. 6, 35249.

Zhang, Y., Kwok-Shing Ng, P., Kucherlapati, M., Chen, F., Liu, Y., Tsang, Y.H., de Velasco, G., Jeong, K.J., Akbani, R., Hadjipanayis, A., et al. (2017). A Pan-Cancer Proteogenomic Atlas of PI3K/AKT/mTOR Pathway Alterations. Cancer Cell 31, 820–832.e3.

