## Supplementary figures and images for "The lipid kinase PI3Kα is required for the cohesion and survival of cancer cells disseminated in serous cavities"

### Suppl Fig S1 and 2

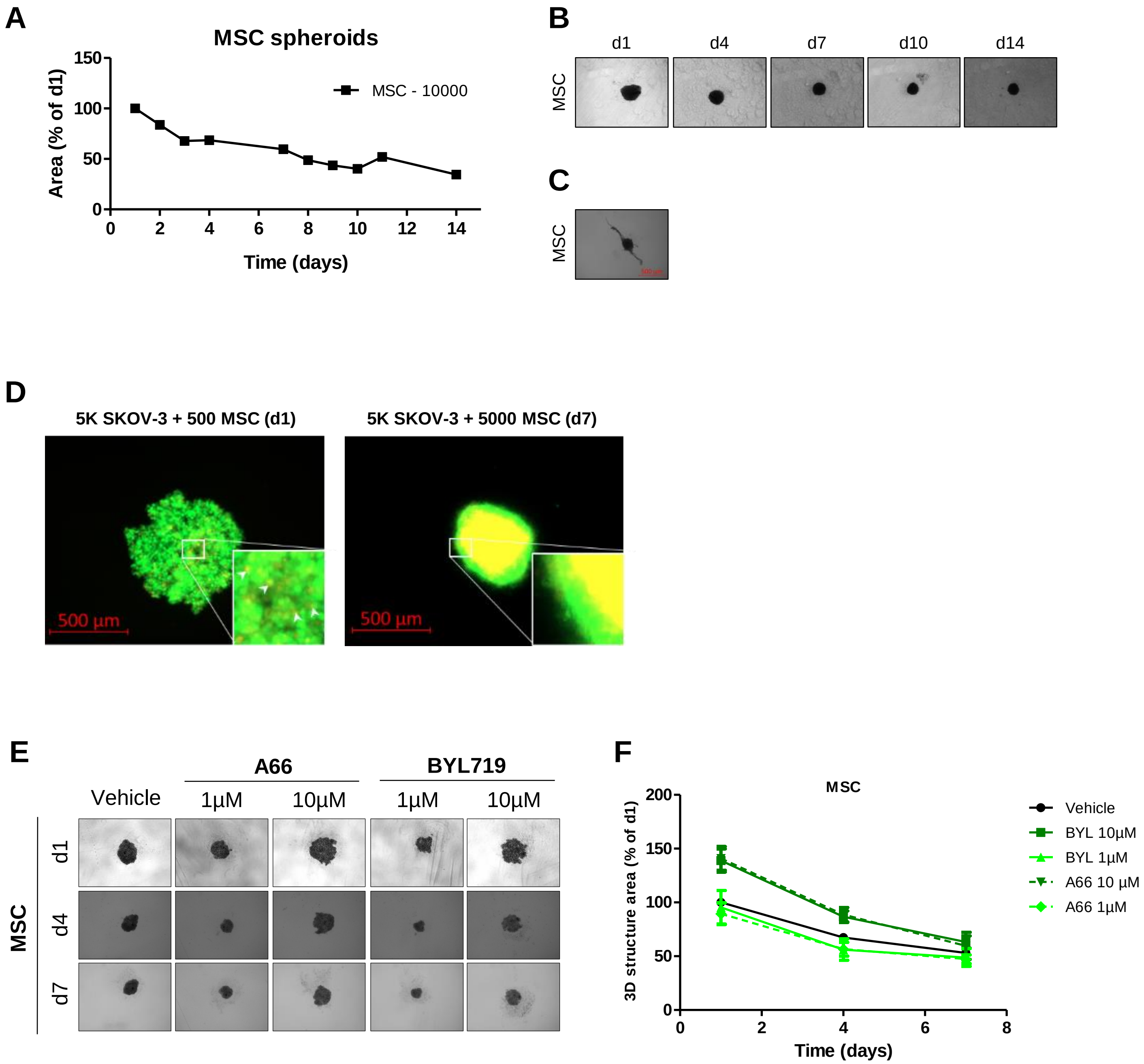


#### Supplementary Figure S1

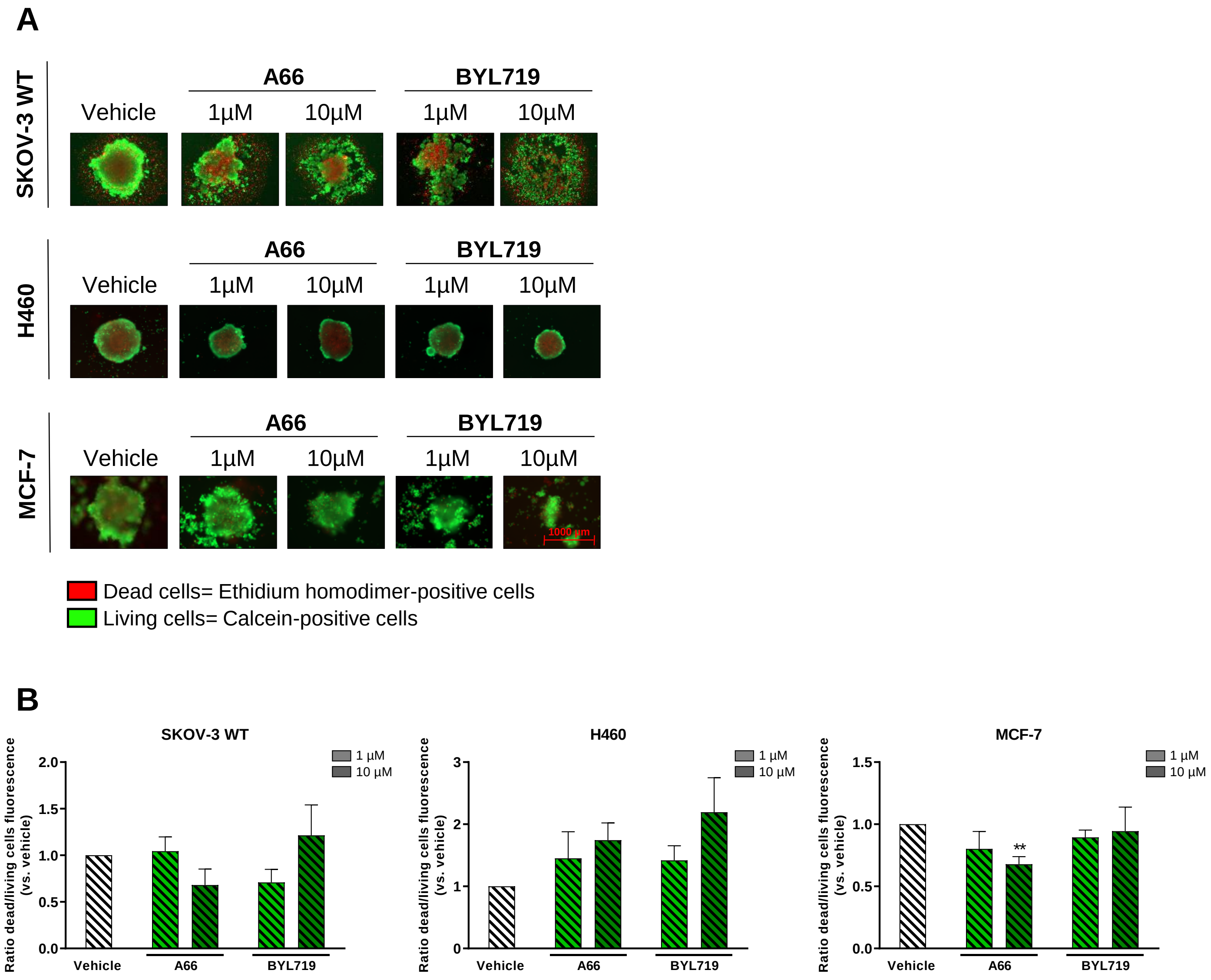


#### Supplementary Figure S2
