## Supplementary material for "The lipid kinase PI3Kα is required for the cohesion and survival of cancer cells disseminated in serous cavities": Suppl Table 1

| **Patient #** | **Age (years)** | **Histology** | **Cohesive Aggregate numbers (per field)** |
| --- | --- | --- | --- |
| 1 | 72 | high-grade serous | 12 |
| 2 | 65 | high-grade serous | 12 |
| 3 | 70 | high-grade serous | 3 |
| 4 | 61 | high-grade serous | 120 |
| 5 | 57 | high-grade serous | 104 |
| 6 | 66 | high-grade serous | 23 |
| 7 | 65 | high-grade serous | 3 |
| 8 | 73 | high-grade serous | 36 |
| 9 | 58 | high-grade serous | 5 |
| 10 | 61 | high-grade serous | 3 |
| 11 | 62 | high-grade serous | 12 |
| 12 | 56 | high-grade serous | 8 |

**Supplementary Table S1**
