## Supplementary material for "The lipid kinase PI3Kα is required for the cohesion and survival of cancer cells disseminated in serous cavities": Suppl Table 2

| **Cel line** | **Cell origin** | ***PIK3CA* mutation** | ***TP53* mutation** |
| --- | --- | --- | --- |
| **OCI-P5x** | Tumor | WT | Y236N |
| **OCI-U1a** |  | WT | V272M |
| **OCI-P8p** |  | WT | S127F |
| **OCI-P2a** |  | WT | R267G |
| **FCI-P2p** |  | WT | R248Q |
| **OCI-P9a1** |  | WT | WT |
| **OCI-P7a** |  | WT | WT |
| **OCI-E3x** |  | WT | WT |
| **OCI-E1p** |  | E545G | WT |
| **OCI-C1p** |  | Q546L, P539R | WT |
| **OCI-C5x** |  | WT | WT |
| **OCI-C4p** |  | WT | 342STOP |
| **OCI-C2p** |  | C420R | WT |
| **Caov-3** |  | WT | Mutated |
| **ES-2** |  | WT | S241F |
| **IGROV-1** |  | Mutated | Mutated |
| **A2780** |  | WT | WT |
| **TOV-112** |  | Mutated | R175H |
| **SKOV-3** | Ascitis | Mutated | WT/Null |
| **59M** |  | WT | WT |
| **EFO-21** |  | WT | Mutated |
| **OV56** |  | WT | Mutated |
| **OV90** |  | WT | S215R |
| **PEO1** |  | WT | Mutated |
| **PEO4** |  | WT | Mutated |
| **PA-1** |  | WT | Mutated |
| **PEO16** |  | WT | WT |
| **PEO14** |  | WT | Mutated |
| **MDAH-2774** |  | Mutated | Mutated |
| **OAW42** |  | Mutated | WT |
| **PEO6** |  | WT | Mutated |
| **OAW28** |  | WT | Mutated |
| **HOC7** |  | WT | Mutated |
| **OVCAR-3** |  | WT | Mutated |
| **PEA2** |  | WT | Mutated |
| **PEO23** |  | WT | Mutated |
| **PEA1** |  | WT | Mutated |
| **COV362** |  | WT | Mutated |
| **COV318** |  | WT | Mutated |
| **OV17R** |  | WT | Mutated |

**A**

**A**

| **Cell origin** | **Number of cell lines  mutated on *PIK3CA* and/or *TP53*** | **Percentage of cell lines  mutated on *PIK3CA* and/*or TP53*** |
| --- | --- | --- |
| Tumour | 13/18 | 72 |
| Ascitis | 19/22 | 86 |

**B**

**Supplementary Table S2**
